# Circulating Immune Cells are Associated with Non-Inflammatory Pain in Rheumatoid Arthritis

**DOI:** 10.64898/2026.03.31.715669

**Authors:** Meghan Mayer, Tyler Therron, Cece Stumpf, Morgan Langereis, Gelis Galarce Lugo, Kathleen Aren, Mary Carns, Jing Song, Cally Mills, Cheol Min Lee, Vanessa Manada De Lobos, Mohammad Daud Khan, Matthew Dapas, Lutfiyya Muhammad, Carla M. Cuda, Yvonne C. Lee, Deborah R. Winter

## Abstract

Over half of patients with rheumatoid arthritis (RA) report clinically meaningful pain, despite treatment with disease-modifying antirheumatic drugs (DMARDs). While joint inflammation is a known cause of pain in patients with rheumatic diseases, emerging data indicate that many patients also suffer from centralized or nociplastic pain. There is a critical unmet need to characterize the altered cellular state that distinguishes patients with centralized pain. In the IMPACT study, 39 RA patients with minimal joint inflammation but varying levels of pain underwent quantitative sensory testing (QST) to assess nociplastic pain, completed patient-reported outcome (PRO) surveys, and provided blood samples for immune profiling.

Supervised and unsupervised analysis of the multi-parameter spectral flow cytometry data identified immune cell populations correlated with nociplastic pain and patient-reported pain intensity. Moreover, analyses of single-cell RNA-seq from a subset of 22 patients revealed differences in cell type proportions and differential expression between the high and low pain groups. These studies provide novel insights into the role of circulating immune cells in altered central nervous system (CNS) pain regulation in adults with RA.

## Introduction

Over the past few decades, the U.S. Food and Drug Administration (FDA) has approved many biologic and targeted synthetic disease-modifying antirheumatic drugs (DMARDs), which have transformed the treatment of rheumatoid arthritis (RA). However, pain continues to be a common and costly problem. In a study of 169 adults with established RA, 47% reported moderate to high levels of pain and fatigue.^1^ Similarly, in a study of 11,995 adults with RA initiating biologic DMARDs, over half continued to report clinically significant pain, despite well-controlled inflammation.^2^

These observations suggest that the experience of pain in RA is influenced by more than just peripheral joint inflammation. One potential contributor is the central nervous system (CNS) and how it regulates pain processing. Studies using quantitative sensory testing (QST) have indicated that adults with RA exhibit widespread pain sensitivity,^3^ resulting in higher pain intensity and lower likelihood of response to DMARDs,^4,5^ Mechanistically, these abnormalities are often referred to in the context of central sensitization. Clinically, they are frequently categorized under the framework of nociplastic pain, a term used to denote chronic pain arising from altered CNS pain processing.

Immune dysregulation may serve as an early trigger for developing and/or maintaining abnormalities in CNS pain processing associated with chronic pain, resistant to DMARD treatment. In humans, this concept is supported by the higher prevalence of fibromyalgia among adults with systemic inflammatory conditions compared to the general population.^6,7^ Studies examining the relationship between measures of inflammation (swollen joint count, erythrocyte sedimentation rate (ESR), C-reactive protein (CRP)) and the development of fibromyalgia have been inconsistent, with some studies showing no relationship and other suggesting weak associations.^8–10^ However, these studies have been limited by the non-specific nature of these assessments. While ESR and CRP are commonly assessed measures of systemic inflammation, they are affected by many different factors (e.g., body mass index, comorbidities) and do not reflect specific immune processes.

There is a body of evidence for peripheral immune cell involvement in the maintenance of chronic pain in animal models. In models of chronic inflammatory arthritis, macrophages are consistently observed in the dorsal root ganglion (DRG) and spinal cord,^11,12^ and in a surgical model of osteoarthritis, Miller at al. demonstrated that macrophage infiltration of the DRG was associated with persistent allodynia.^13^ T cells also infiltrate the DRB and spinal cord and play a role in the maintenance of pain in rodent models of neuropathic pain.^14,15^ Together, these studies provide the scientific premise for examining the relationship between peripheral immune cells and pain in adults with RA.

Here, we present initial results from the IMPACT study where we recruited 39 adults with RA across a range of self-reported pain intensity. We excluded patients with more than minimal joint inflammation to limit the impact of peripheral pain due to joint inflammation. Each patient completed patient-reported outcomes (PRO) surveys, gave blood, and underwent QST. PBMCs from all patients were processed for multiparameter spectral flow cytometry (flow). Canonical populations were gated and tested for associations with pain intensity and related pain measures. Flow data was also used for unsupervised analysis to identify *de* novo immune cell populations and calculate correlation with various QST and PRO outcomes. Finally, hashed single-cell RNA-seq (scRNA-seq) libraries were generated for 22 samples categorized by high and low pain intensity. These groups were compared for differences in cell composition and transcriptional signature. Our results indicate associations between circulating immune cell populations and pain outcomes.

## Methods

### Patient Recruitment

The IMPACT study was a cross-sectional, observational study examining the relationship between peripheral blood mononuclear cells (PBMCs) and nociplastic pain in adults with RA. Participants were recruited from a single academic center between November 2022 and December 2024. Eligibility criteria included: 1) meeting either 1987 or 2010 American College of Rheumatology (ACR)/ European Alliance of Associations for Rheumatology (EULAR) Criteria for RA,^16^ 2) on a DMARD for RA, and 3) swollen joint count ≤ 1. Participants were excluded if they had received an intramuscular steroid injection within four weeks of the study visit or were taking corticosteroids at doses > the equivalent of prednisone 10 mg daily, scheduled opioids, or central-acting pain medications (e.g., tricyclic antidepressants, selective serotonin norepinephrine reuptake inhibitors, gabapentin, pregabalin). Participants were also excluded if they were pregnant or had clinically significant peripheral neuropathy, peripheral vascular disease, or another systemic autoimmune disease. Ethics approval was obtained from the Institutional Review Board of Northwestern University. All participants provided written informed consent.

### Clinical Variables and Patient-Reported Metrics

Demographic characteristics were obtained by self-report. A trained examiner performed tender and swollen counts of 28 joints. Patient reported pain intensity was assessed using a 0-10 numeric rating scale (NRS). The extent of fibromyalgia symptoms was assessed using the Fibromyalgia Survey Questionnaire (FSQ), which is comprised of the Widespread Pain Index (WPI) and Symptom Severity Scale (SSS).^16,17^ Cognitive function was assessed using a single item within the SSS, asking participants to rate trouble thinking or remembering on a 0-3 scale, with 0 being no problem and 3 being severe problems. Sleep disturbance and fatigue were assessed using four-item Patient-Reported Outcomes Measurement Information System (PROMIS) computerized adaptive tests.^18,19^

### Quantitative Sensory Testing (QST)

To characterize nociplastic pain, participants completed QST to measure pressure pain thresholds (PPTs), using previously published methods.^4^ PPTs were measured at bilateral trapezius muscles (non-joint site) using a Force 10 FDX algometer (Wagner). Pressure was increased at a rate of 0.5 kgf/sec until the participant first reported pain. Three separate measurements were collected on each side, with at least 30 seconds between tests. The results were averaged to obtain a single measure of PPT at the trapezius. In adults with RA, lower PPTs at the trapezius indicate central sensitization, consistent with nociplastic pain.^20^.

### Processing of PBMCs

Blood was collected from patients and stored in 3 EDTA tubes overnight while being rocked. Samples were diluted 1:1 with DPBS (no calcium, no magnesium) and split over three 50mL Greiner Leucosep tubes pre-loaded with 15mL Histopaque 1077. Samples were centrifuged at 1000g for 10 minutes at room temperature with no brake. The buffy coats were removed, combined into one tube, diluted up to 50mL with DPBS and centrifuged at 350g for 10 minutes at room temperature. The pellet was resuspended in an additional 50mL of DPBS and centrifuged under the same conditions. Red blood cell lysis was performed by resuspending the pellet in 5mL of Gibco ACK lysing buffer and incubating for 5 minutes at room temperature.

Samples were diluted with 4C AutoMACS Running Buffer (MACS) and centrifuged at 350g for 10 minutes at 4C. The pellets were resuspended in MACS, filtered, counted and again centrifuged at 350g for 10 minutes at 4C. Cryostor CS10 was added to give a concentration of 5 million live cells/mL and samples were aliquoted at 1mL into cryovials. The samples were frozen using a Mr Frosty Freezing container and then transferred from -80C to liquid nitrogen after 1 week for longer storage.

### Multiparameter flow cytometry

Samples from 39 patients were split into 3 batches of 13-14 samples taking care to keep demographics, PRO and QST metrics equal between batches (patient IMP29 was repeated daily as a control). For each batch, frozen samples (1 tube of 5 million cells per sample) were removed from liquid nitrogen and transferred to a 37C water bath until thawed. In groups of 4, thawing media (0.5% BSA, 1X Glutamax and 10mM HEPES in RPMI) was added slowly in an equal volume to the sample and then transferred over a 20um filter into a new tube while on ice. The filters were rinsed with an additional 8mL of thawing media, and the samples were centrifuged at 350g for 10 min at 4C (all subsequent centrifugations use these conditions). The pellets were washed with cold HBSS and aliquots were taken for counting. After centrifuging again, the pellets were resuspended in HBSS containing eFlour 506 Fixable Viabilty dye and incubated at room temperature for 15 minutes. Samples were diluted with cold MACS and centrifuged to stop the staining, then rinsed an additional time with MACS. Pellets were resuspended in FC block for 20 minutes at 4C, then antibody cocktail was added for an additional 30 minutes. Samples were diluted with MACS, centrifuged and rinsed one additional time with MACS. PFA was added (2%), and samples were incubated at 4C for 20 minutes to fix. Following fixation, samples were rinsed with MACS and stored at 4C overnight. FMOs were thawed and stained in tandem, using samples from control patients. The following day, samples were analyzed using the BD FACSymphony A5.2 Spectral Analyzer.

### Unsupervised analysis of flow data

The samples for each batch were compensated separately, and initial gating was performed to select live, CD45+ cells and exclude granulocytes. Compensated samples were exported at 70,000 cells per sample then concatenated together. Cycombine was performed on the concatenated file to normalize batch variation. Each fluorescent parameter was adjusted to ensure cells were on scale, and the data was visualized using UMAP (Euclidean distance metric, 15 nearest neighbors, 0.5 min distance, all compensated parameters except CD45 BB515 and L/D BV480). FlowSOM was used to generate clusters using the same compensated parameters and 20 clusters as input. Clusters were evaluated by visualizing the MFI of each fluorescent parameter per cluster and comparing distribution to known populations.

### Generation of Single-Cell Libraries

A subset of 22 patients exhibiting high and low pain were chosen for single cell analysis. Samples were split into 3 batches with the first batch consisting of 3 high and 3 low pain samples, the second and third batches consisting of 4 high and 4 low pain samples each. All samples were thawed in the same manner as above.

For batch 1, following the first centrifugation, samples were resuspended in MACS, counted and 1 million total cells were pulled forward for staining and centrifuged. The pellets were resuspended in FC block, incubated for 20 minutes at 4C, then antibody cocktail containing anti-CD15 AF700 and a unique TotalSeq-C anti-human hashtag and CD45 fluorophore for 30 min at 4C. The samples were rinsed twice with 5mL of MACS, then resuspended in a DAPI solution and pooled. The FACSymphony S6 Spectral Sorter was utilized 6-way sort 50000 live, CD45+, CD15- cells for each sample. The samples were pooled, filtered, centrifuged and resuspended in loading buffer (0.04%BSA, 1X Glutamax, 10mM HEPES in RPMI). Following counting, 24000 cells were loaded into Chromium Next GEM Chip K using aa Chromium iX controller. cDNA and GEX, ADT (Antibody-Derived Tags), and TCR libraries were made following the manufacture directions for Next GEM 5’ Single cell kits (10X Genomics). GEX and TCR libraries were prepared using the Chromium Connect. Quality of cDNA and libraries was assessed using the Aligent Tapestation and Invitrogen Qubit. All libraries for batch 1 were pooled and sequenced on the NovaSeq X at a targeted depth of 800M reads PE100.

For batches 2 and 3, 5 millions cells were FC blocked for 20 minutes at 4C. Each of the 8 samples per batch was stained with anti-CD15 AF700 and a unique TotalSeq-C anti-human hashtag but split into 2 groups of 4 unique CD45 fluorophores. The samples were rinsed twice with MACS then counted, and the 2 groups of 4 samples were pooled at 150000 cells/sample. Samples were stained with Antibody-Derived Tags (ADTs) from the TotalSeq-C universal cocktail at for 30 minutes at 4 C. Following a rinse with MACS, samples were resuspended in a DAPI solution. Using the FACSARIA sorter, the 2 groups of 4 per batch were sorted for 25000 live, CD45+, CD15- cells per sample. The samples from both sorts were pooled and, counted and 32000 cells were loaded into 2 lanes of a Next GEM Chip K using the Chromium Connect (10X Genomics). cDNA and GEX, ADT and TCR libraries were prepared, and QC performed as for batch 1. All libraries for batches 2 and 3 were pooled and sequenced on the NovaSeq X Plus over 2 lanes for a targeted depth of 3 billion reads.

### Processing & Analysis of Single-cell Libraries

The single-cell libraries were aligned to GRCh38 human transcriptome and processed using the CellRanger multi pipeline command, version 7.1.0 with default parameters. The Universal Human TotalSeq C antibody panel from BioLegend served as surface marker reference for batches 2 and 3 (WHL 2-5). Demultiplexing of the cell hashing was done using HTODemux from Seurat v5, with HTO assay count data normalized using CLR (margin = 1). Cells were filtered with thresholds of nCount_RNA >= 2000 and nCount_Ribo >= 150. IMP12 was excluded because of low-quality concerns. Technical replicates across pools were combined into one sample for succeeding analyses. Default parameters from the NormalizeData function in the Seurat package were applied to the RNA assay. Highly variable genes were identified using the FindVariableGenes function, followed by data scaling with ScaleData.

Neighbor finding, clustering, and UMAP generation were based on principal components (PCs) 1-9 and 11-12. The tenth PC was excluded due to the absence of meaningful variance within it. Neighbor graph construction and UMAP generation were executed with default parameters. Cells were clustered using the Louvain algorithm with a resolution of 0.3. Differential composition analysis was performed using the MASC R package. Differential expression (DE) analysis was performed on pseudobulk cell types using the DESeq2 Bioconductor package. Pseudobulk count matrices were generated by the summation of counts across cells from the same donor within each cell type. Ribosomal, mitochondrial, and Y chromosome genes were excluded prior to DE analysis. DESeq2 models were fitted to Pain Group identity with age and sex added as covariates to the design matrix; the reference condition was “Low Pain”. Expressed genes with a minimum of 10 counts in at least 3 of the pseudobulk samples were included in the DE analysis. Gene set enrichment analysis (GSEA) was performed using the fgsea R package on ranked genes results from the DE analysis. Rankings were determined from the Wald statistic as part of the DESeq2 output data tables. Following the input of this ranked gene vector into fgsea, Hallmark gene sets were gathered from MSigDB using the msigdbr R package for *Homo Sapiens*. Pathways with less than 15 genes or more than 500 genes were filtered out from downstream analyses. Normalized enrichment scores (NES) were calculated for each pathway and those with a nominal p-value less than 0.05 were considered significant.

### Statistical Analysis

Associations between cell population proportions and outcome metrics were assessed by Pearson correlation coefficients. Correlations above 0.5 were considered strong, 0.25-0.5 moderate, and 0.1-0.25 weak.

## Results

### Testing Associations between Canonical Immune Cell Populations and Pain

We obtained blood and assessed pain levels in 39 adults with RA exhibiting swollen joint count ≤ 1 (**Figure 1A**). The demographics, disease parameters, and pain levels of these patients are summarized in **Table 1**. We used a multiparameter spectral flow cytometer to immunophenotype PBMCs from each individual. After excluding any contaminating granulocytes, we gated on 25 subpopulations of CD4 T cells, CD8 T cells, B cells, Natural Killer (T) cells (NK/NKT), Monocytes, and Dendritic Cells (DCs) (**Supp Figure 1A**). On average, we found patients exhibited 24.7% CD4 T cells, 12.2% CD8 T cells, 12.3% B cells, 15.1% NK/NKT cells, 24.4% monocytes, and 2.07% DCs (**Figure 1B**). We found a moderate positive correlation between total monocytes and pain intensity (**Figure 1C)**. We also found weak negative correlations between CD4+ T cells and CD8+ T cells with pain intensity (**Figure 1C)**. For the average PPT at the trapezius, a QST measure that decreases with nociplastic pain, we found a weak positive correlation with total B cells and DCs (**Figure 1D**). At the subpopulation level, we observed moderate negative correlations of CD4+ Naïve T cells and CD8+ naïve T cells and moderate positive correlation of classical monocytes (CM) with pain intensity, suggesting these subpopulations drive the earlier cell type level correlations (**Supp Figure 1B).** We found that cDC1 exhibited the highest correlation with PPT. Interestingly, we rarely observe a correlation in one subpopulations with both pain metrics; in our study, pain intensity and average PPT were only weakly correlated (**Supp Figure 1C**). Taken together, we observe higher correlations between known circulating immune populations and patient reported pain intensity than with the QST measure of nociplastic pain.

**Figure 1.**
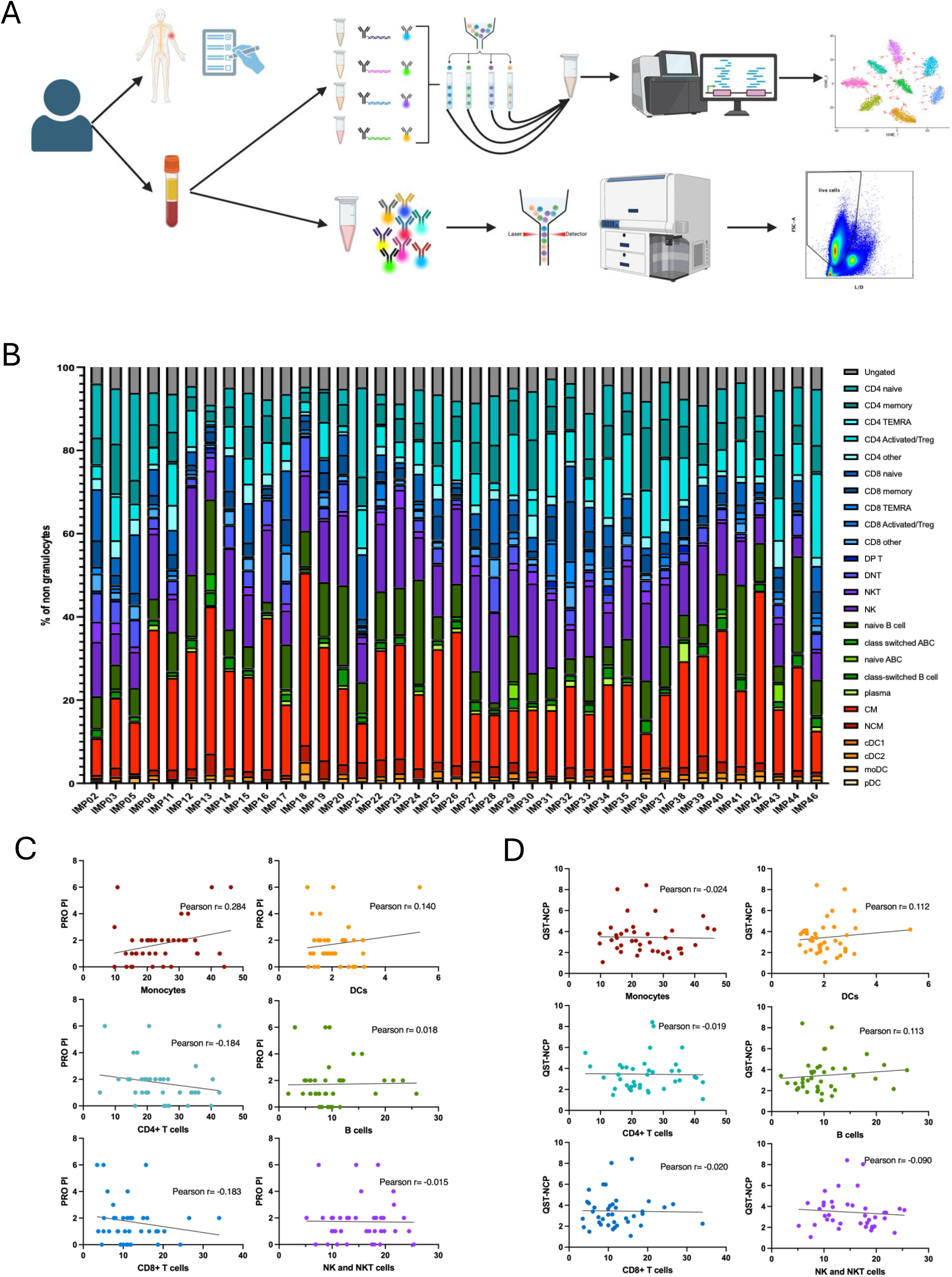
Associations between known immune cell populations and pain. (**A)** Overview of IMPACT study design. **(B)** Known cell populations per patient shown as percent of non-granulocyte PBMCs. DP T= double positive (CD4+/CD8+) T, DN T= double negative (CD4-/CD8-) T cells, NKT= Nature Killer-like T cells, NK= Natural killer cells, ABC= Age-associated B cells, CM= Classical monocytes, NCM= Non classical monocytes, cDC1 and cDC2= conventional DC 1 and 2, moDC= monocyte-derived DC, pDC= plasmacytoid DC. **(C-D)** Scatterplots showing correlation of known cell population proportions with patient reported numeric rating scale for pain intensity (PRO PI) (C) and nociplastic pain as measured by quantitative sensory testing of pressure pain threshold at trapezius (QST-NCP) (D).

**Table 1.** Demographic and clinical characteristics (N = 39).

| **Table 1. Demographic and clinical characteristics (N = 39).** |  |
| --- | --- |
| **Characteristic** | **Mean (SD) or Number (%)** |
| Age, years | 54 (16) |
| Female | 32 (82%) |
| Seropositivity | 31 (79%) |
| Disease duration, years | 11 (11) |
| Clinical Disease Activity Index (CDAI) | 3.49 (3.34) |
| Fibromyalgia Survey Questionnaire (FSQ) | 4.82 (2.93) |
| Widespread Pain Index (WPI) | 1.92 (1.97) |
| Symptom Severity Scale (SSS) | 2.90 (2.20) |
| Pain Intensity NRS, 0-10 | 1.7 (1.6) |
| PROMIS Sleep Disturbance | 49 (9) |
| PROMIS Fatigue | 48 (8) |
| Trapezius PPT | 3.45 (1.62) |

### Unsupervised Analysis of Immune Cells Reveals Correlations with Various Outcomes

To identify novel populations among our samples that may correlate with pain and related outcomes, we performed an unsupervised analysis of the flow data from a down-sampled cell subset of each patients (**Supp Figure 2A)**. We defined 20 clusters with distinct surface marker profiles that were conserved across batches (**Figure 2A, Supp Figure 2B-C)**. The majority of these aligned cleanly with high level annotations including CD4+ T cells, CD8+ T cells, NKTs, B Cells, DCs, or monocytes (**Figure 2B, Supp Figure 2D-E**); however, there were 4 clusters that were previously undefined. Next, we calculated the correlation between each of these clusters and patient reported outcomes for pain, cognition, and sleep (**Figure 2D**). We found that Mono1 CD89^high^ was moderately positive correlated with pain intensity which is consistent with classical monocytes in **Supp. Figure 1**. Surprisingly, Mono1 is also moderately positive correlated with the Widespread Pain Index (WPI) score, despite the fact that these two metrics are not correlated with each other (**Supp Figure 2F**). Mono2 CD88^high^, which was not correlated with Mono1 (**Supp Figure 2G**), also exhibited moderately positive correlation with WPI, but moderately negative correlation with the trouble thinking, a metric that increases with worsened cognition. Of T cells, we observe a moderate negative correlation of CD4 T2 CD45RA+ and CD8 T2 CD45RA+ with pain intensity which validates earlier results with naïve CD4+ and CD8+ T cells. We observed the highest negative correlation between Other4 and pain intensity. Other4 is a CD3/CD15- cluster that does not align well with any known populations and comprises approximately 0.5% of the total cells. While DC1 CD141+, which overlaps cDC1, exhibits weaker correlation with nociplastic pain than its known counterpart and no correlation with pain intensity, we observed moderately negative correlation with both sleep-related metrics (PROMIS Sleep Disturbance and Fatigue). We found the highest correlation involving nociplastic pain is with B2 CD11C+, containing both naïve and class-switched age-associated B cells (ABC). CD11C^high^ B cells have previously been associated with age and autoimmunity.^21^ DC3 exhibit the next highest correlation with PPT which is interesting because, like pDCs and naïve ABCs in the prior analysis, DC3 is one of the few clusters correlated with pain intensity in same trend. In addition, we found that DC3 hasthe highest negative correlation with the PRO for sleep disturbance. These results suggest that the immunophenotype of circulating immune cells is reflective of various aspects of pain status.

**Figure 2.**
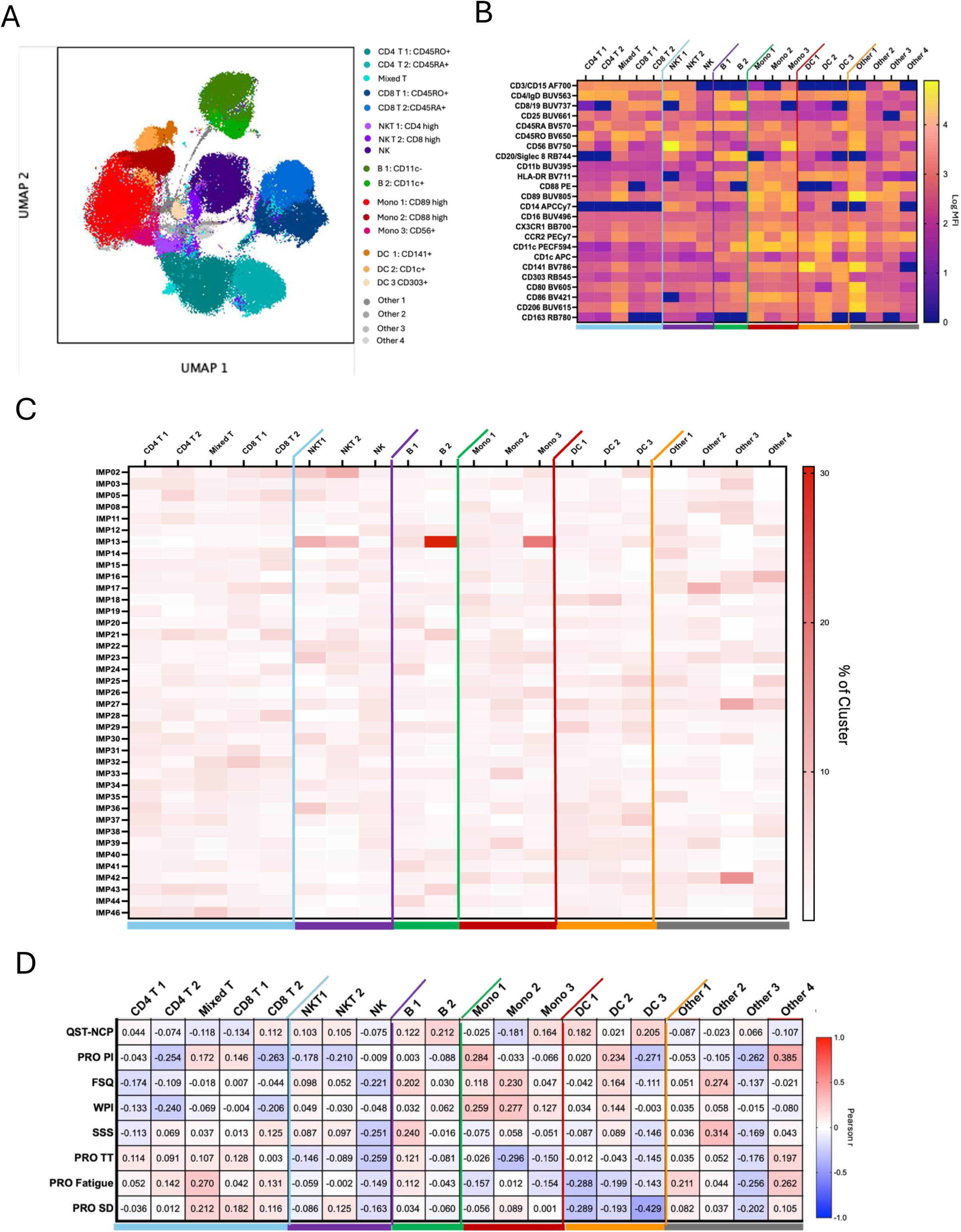
Unsupervised analysis of multiparameter spectral flow cytometry. (**A**). UMAP of concatenated samples annotated with unsupervised clusters. (**B**) Fluorescence intensity of unsupervised cluster shown as log of normalized MFI (Geometric mean). (**C**) Percent of each cluster originating from each patient. (**D**) Pearson correlation between unsupervised cluster proportions and pain-related outcomes. QST-NCP= Quantitative Sensory Testing for Nociplastic Pain by pressure pain threshold at trapezius, PRO PI= Patient-Reported NRS for Pain Intensity, FSQ= Fibromyalgia Survey Questionnaire, WPI= Widespread Pain Index, SSS= Symptom Severity Scale, PRO TT= PROMIS Trouble Thinking or Remembering, PRO Fatigue= PROMIS Fatigue, PRO SD= PROMIS Sleep Disturbance.

### Comparison of Transcriptionally Defined Immune Cell Populations in RA Patients with Low Pain

To investigate whether there were transcriptional states of circulating immune cells that varied with pain, we performed scRNA-seq on 36,594 PBMCs from a subset of 22 IMPACT patients categorized into high and low pain intensity groups **(Supp Figure 3A**). We defined 11 transcriptional distinct clusters across groups including canonical populations of Naïve CD4+ T cells, Memory CD4+ T cells, CD8+ T cells, NK/NKT cells, B cells, CM, NCM, cDC, and pDCs that expressed known gene markers and canonical surface markers (**Figure 3A-C, Supp Figure 3B-C**). Annotations were confirmed by surface marker expression as measured by ADT (**Figure 3D**). The proportions of these populations by sample largely agree with those quantified by flow cytometry (**Supp Figure 3D-E).** Interestingly, our clustering approach also identified distinct T cell subpopulations exhibiting interferon-responsive (IFN) genes. We found that these IFN subpopulations were largely populated by cells in the low pain intensity group, but the trend was driven by only a few patients. On the other hand, NK/NKT cells were significantly increased in the high pain intensity group compared with the low pain intensity group (**Figure 3E).** As in the flow data, we tend to see fewer T cells and more monocytes with higher pain intensity. The observed changed in composition indicate that there may be underlying transcriptional differences between pain intensity groups.

**Figure 3.**
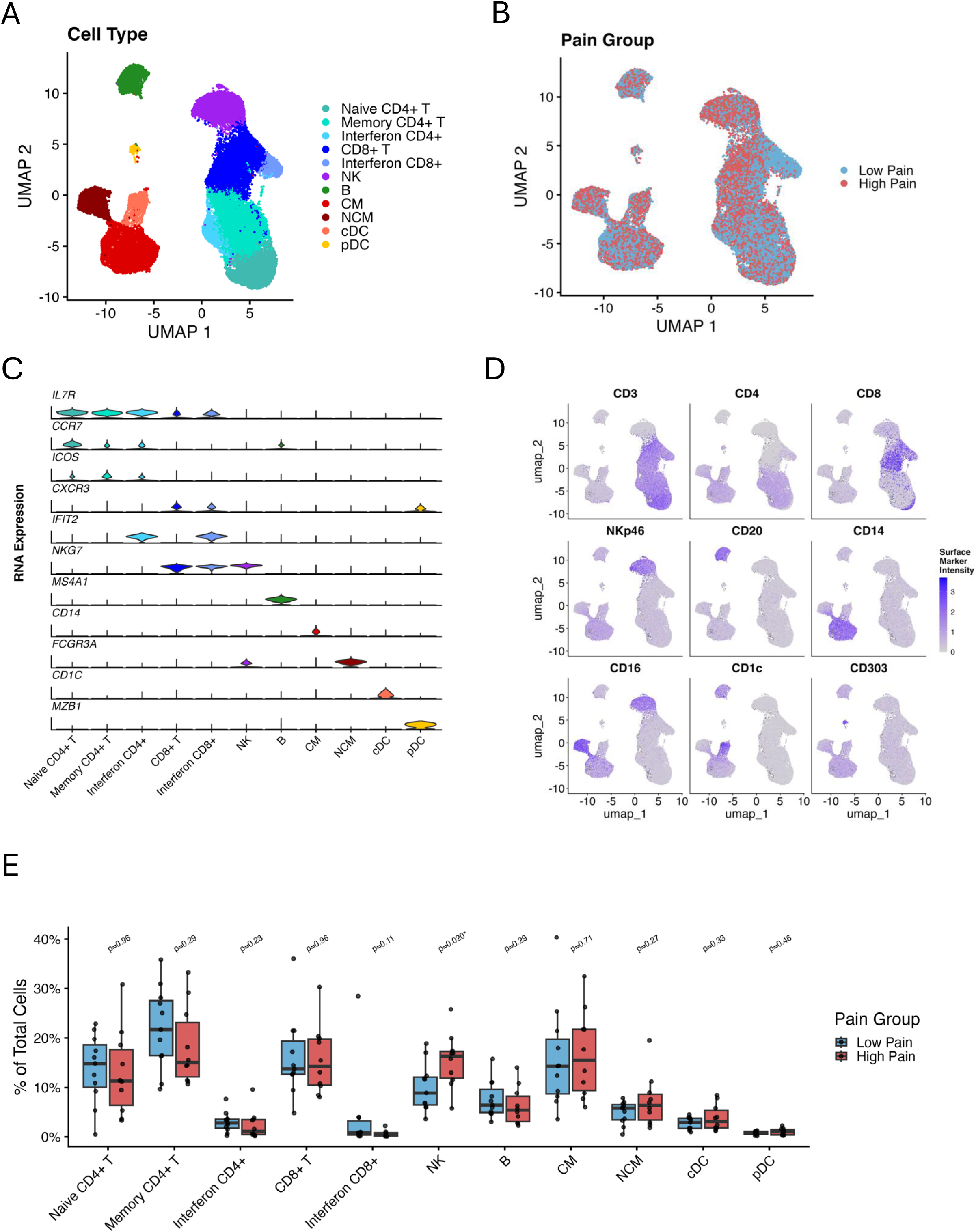
Single-cell RNA-seq of PBMCs from low and high pain groups. (A) UMAP annotated by cell type. (B) UMAP annotated by pain group. (C) Violin plots showing RNA expression of known gene markers. (D) Feature plots showing canonical surface marker intensity as measured by ADT. (E) Box and whisker plot of cell population proportions split by pain group. P-values were calculated by MASC.

### Differential Expression between High and Low Pain Intensity Groups

Next, we performed differential expression on pseudobulk expression between high and low pain intensity groups in major cell types (**Supp Figure 4A**). We found the greatest number of differential genes (191) in CD8+ T cells (**Figure 4A, Supp Figure 4B**). Many of these (23) were shared with either CD4+ T cells or NK/NKT (**Supp Figure 4C**). Monocytes also exhibited many differential genes (135) and 8 of these were shared with DCs including IRAK3 (high), RNF44B (high), and HLA-DQB2 (low) (**Supp Figure 4D)**. Interferon-responsive genes were associated with low pain intensity across cell types, including *IFIT3, OAS2*, and *OASL* which were significantly differential in at least CD8+ T cells (**Figure 4C**). Genes in the *Angionesis* and *Myogenesis* pathways, including *PF4*, OLR1, and *MEF2D* were more highly expressed in monocytes from the low pain intensity group with some overlap in B cells. In the high pain intensity group, CD8+ T cells exhibited increased expression of genes from the TNFA Signaling, Apoptosis, and Inflammatory Response, such as *TNFAIP3, SMAD3, and CEBPD*. Interesting, the same pathways were associated with low pain intensity in NK cells. On the other hand, high pain NK cells highly expressed genes involved in WNT/Beta Catenin Signalling, including *JAG1, DLL1, and AXIN1.* In summary, we observed significant pain-associated differences in the transcriptional profile of circulating immune cells.

**Figure 4.**
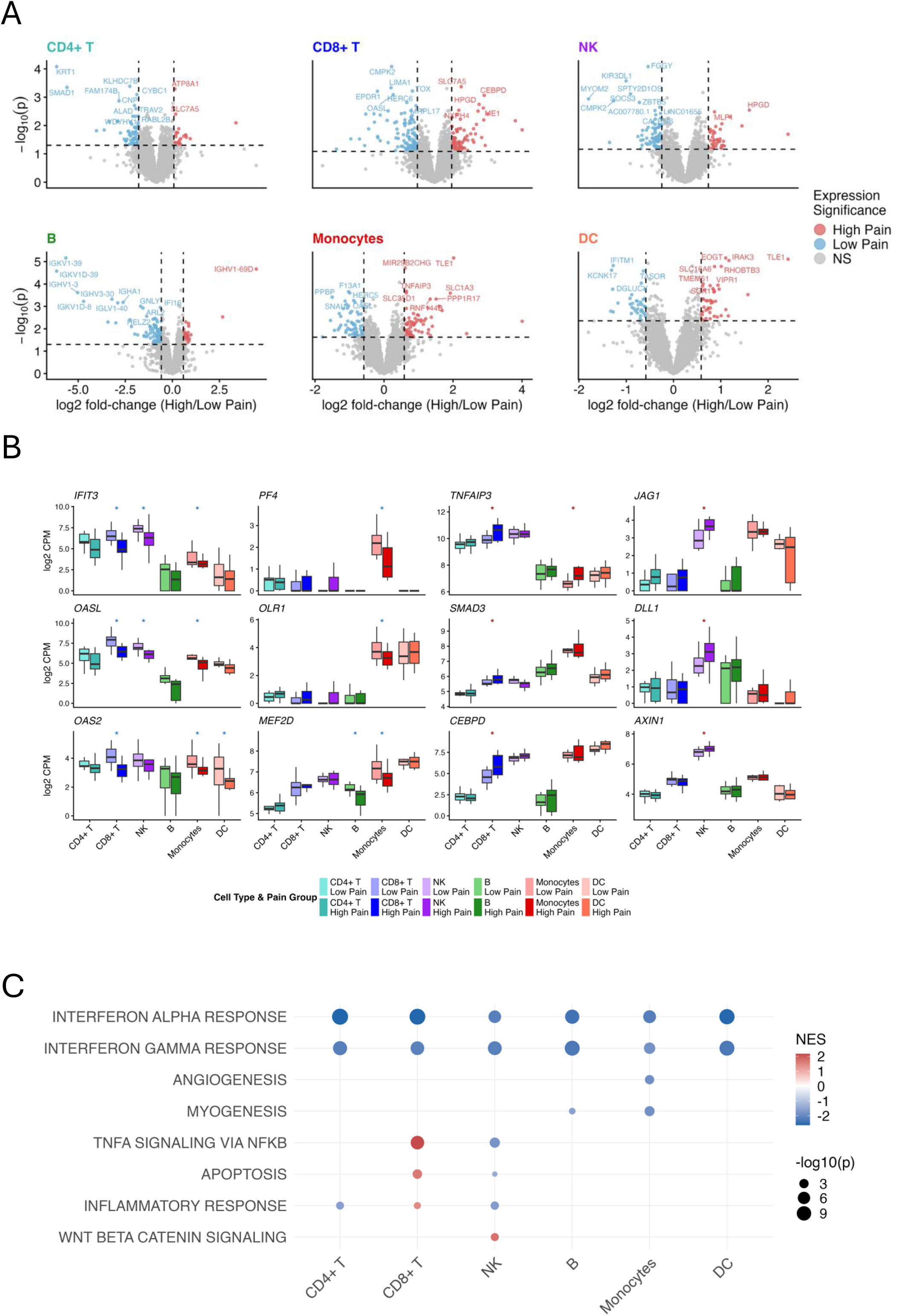
Pseudobulk differential expression between low and high pain groups. (A) Volcano plots of differentially expressed genes (DEGs) by cell type. Color indicates DEGs (DEGs, FC>|1.5|, p<0.05) associated with low (blue) and high (red) pain. (B) Box and whisker plots of pseudobulk expression of select genes across cell types. Asterisks indicate significant association (DEseq p < 0.05) with low (blue) or high (red) pain by cell type. (C) Bubble plot of normalized enrichment score (NES, color) and -log10 p-value (size) of select Hallmark pathways enriched in at least one cell type (p <0.05). NES and p-values calculated GSEA.

## Discussion

In this study, we investigate associations between pain metrics and circulating immune cells in adults with RA. We find weak to moderate correlations between several known immune cell populations and pain intensity, as well as PPT trapezius, a QST metric for nociplastic pain. Unsupervised annotation of immune cells largely recapitulated these findings and revealed additional correlations with related outcomes. Analysis of single-cell RNA-seq revealed a significant increase in NK cells in the high pain intensity group. Finally, we found CD8+T cells exhibited the most differential genes and including an interferon-responsive signature associated with low pain that was shared across cell types. Taken together, these results support a role for circulating immune cells in pain in RA.

Across both continuous and group-based analyses, higher pain intensity was associated with lower T cell proportions and higher monocyte proportions. Prior scRNA-seq studies have varied in reports of changes to immune cell composition between RA and controls.^22–24^ A study by Binvignat et al. stratified their patient cohort by a composite disease activity score (DAS28-CRP) and found increased CD4+ memory T cells and non-classical monocytes in patients with high disease activity.^22^ To our knowledge, no studies have specifically examined the relationship between scRNA-seq and pain in adults with RA. We designed our study to exclude participants with more than one swollen joint to isolate pain from peripheral joint inflammation. Thus, it is not surprising that our results would not agree with prior reports.

The single-cell analyses revealed immune cell differences associated with pain intensity beyond those observed in the flow analysis. Surprisingly, we found a larger proportion of NK cells in the group with high pain intensity, despite lack of correlation with known and unsupervised NK populations. It may be explained by differences in the definition of the NK cells which is largely defined by negative gating in flow. It is also possible that it is related to the specific subset of patients who were included in the scRNA-seq studies. Previous studies have implicated NK cells in both promoting and controlling RA,^25^ and NK cells are thought to have a protective role in neuropathic pain conditions.^26^ In addition, transcriptionally based clustering defined distinct T cell subsets expressing interferon-responsive genes. A similar CD4+ Tcell population was previously observed in RA patients and slightly, but not significantly, increased in RA patients with high disease activity. ^22^ Moreover, interferon response pathways were among the most enriched with genes associated with the low pain intensity group in all cell types. Recent studies suggest that interferon, especially Type 1 (A/B) may play an analgesic role in nociplastic pain.^27^ Additional research is needed to clarify the role of both NK cells and interferon in RA-associated pain.

RA patients often experience pain, sleep disturbance, fatigue, and psychosocial distress in tandem.^1^ While many of the pain-specific and pain-related metrics we investigated were intercorrelated, their relationships with immune cell type proportions were often discordant. This discordance may provide an opportunity for patients to be stratified into distinct outcome groups. For example, the pDC-like unsupervised cluster (DC3) was associated with lower sleep disturbance but exhibited increasing trends with both patient-reported pain intensity and QST-measured nociplastic pain (manifested by lower trapezius PPTs). This pattern suggests that pDC numbers may help distinguish a subgroup of patients whose pain levels are closely linked to sleep quality. Similarly, some immune cell types were correlated with both pain intensity and WPI, while others were not. Cell types that correlate with both pain intensity and WPI may be more likely to reflect nociplastic pain, whereas those correlating with pain intensity alone may reflect other types of pain. Further study on larger cohorts will be needed to clarify the existence and clinical relevance of these subgroups.

There are several limitations to this study related to experimental design and the technologies being used. To reduce batch effect of individual samples, we froze all PBMCs within one day of collection so that they could be processed at a future date in large batches. However, samples are likely to have experienced cell death during the freezing/defrosting process that may have influenced cell proportions. Furthermore, while we custom-designed our multispectral flow antibody panel to cover as many cell types and activation states as possible, we were not able to include every informative marker. This limits both our ability to gate known populations, especially NK cells as described above, and the efficacy of the unsupervised clustering on flow data. The scRNA-seq data does not suffer from the same limitation since it provides genome-wide transcriptional data. The issue with scRNA-seq technology is that it represents a sampling of the total cells and cannot be reliably used for cell quantification. Finally, all the analytical approaches suffer from the limited samples size which hampers our ability to infer significance.

In conclusion, the analysis presented here demonstrate associations between circulating immune cells and pain levels in adults with RA. These results support a multifactorial role for various immune cell populations in the development and maintenance of nociplastic pain. A model combining these different factors may be useful as a biomarker marker of prediction of pain in RA. Further studies with increased cohorts and longitudinal sampling would better capture individual variability and changes over time. In addition, studies including medications or behavioral interventions designed to improve pain would demonstrate the use of this approach in predictive models. Our ultimate goal is to gain a better understanding of pain mechanisms in RA to improve development and assessment of therapeutic interventions.

---

**Supp. Figure 1.**
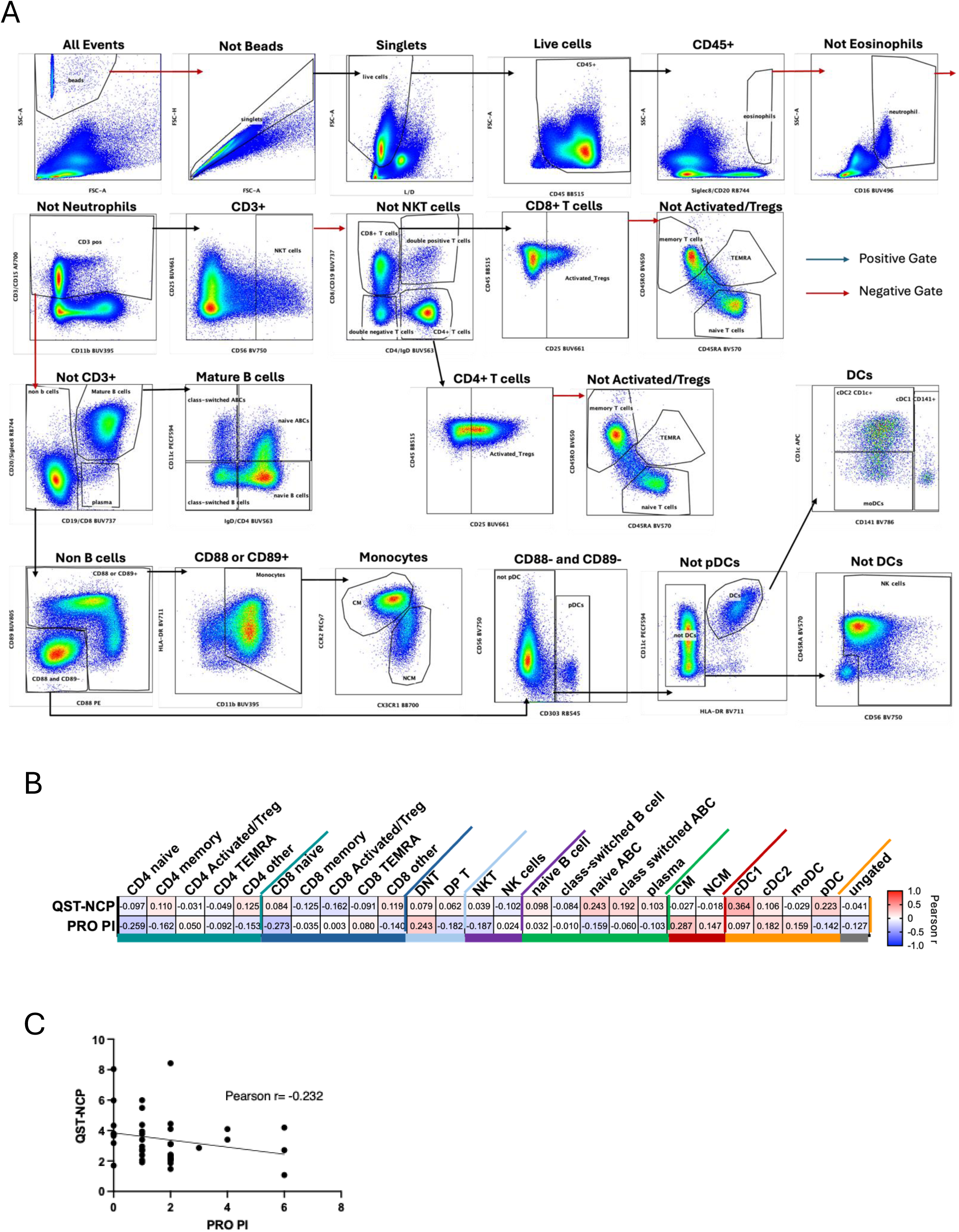
Flow cytometry gating schema and pain metric correlations. (**A**) Gating strategy for known cell populations. Black arrows represent positive gating and red arrows represent negative gating. (**B**) Pearson Correlation of known population proportions with nociplastic pain as measured by quantitative sensory testing of pressure pain threshold at trapezius (QST-NCP) and patient-reported numeric rating scale for pain intensity (PRO-PI). (**C**) Scatterplot showing the correlation between QST-NCP and PRO-PI.

**Supp. Figure 2.**
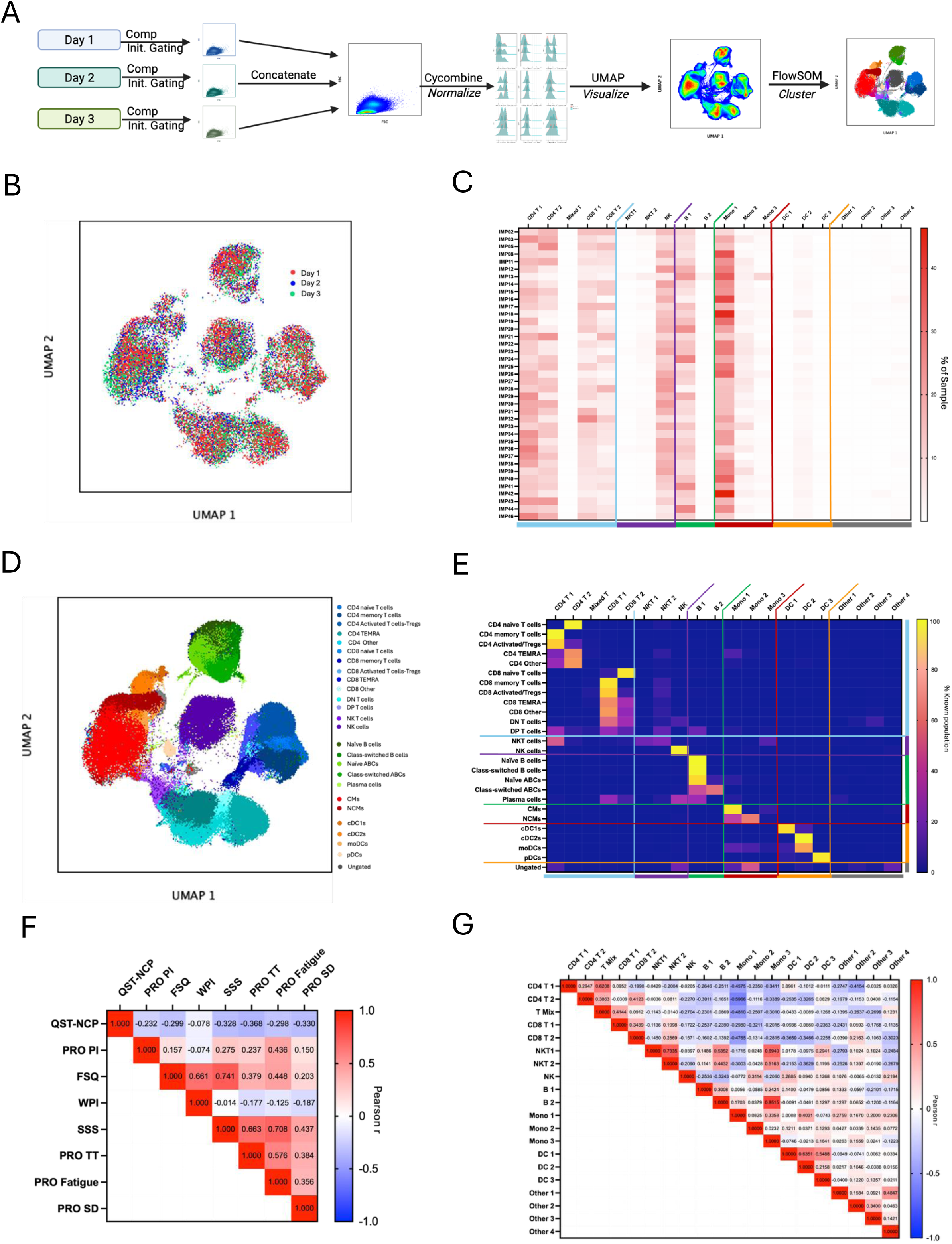
Additional unsupervised flow analysis. **(A)** Overview of workflow for unsupervised analysis of multiparameter spectral flow cytometry. (**B**) UMAP annotated by sample batch (**C**) Percent of each patient sample by unsupervised cluster annotations. (**D**) UMAP annotated by known cell populations. (**E**). Heatmap showing the % of each known cell population by unsupervised cluster_annotation. (**F**) Pearson correlation between pain-related outcomes (**G**) Pearson correlation between unsupervised cluster proportions. QST-NCP= Quantitative Sensory Testing for Nociplastic Pain by pressure pain threshold at trapezius, PRO PI= Patient-Reported NRS for Pain Intensity, FSQ= Fibromyalgia Survey Questionnaire, WPI= Widespread Pain Index, SSS= Symptom Severity Scale, PRO TT= PROMIS Trouble Thinking or Remembering, PRO Fatigue= PROMIS Fatigue, PRO SD= PROMIS Sleep Disturbance.

**Supp. Figure 3.**
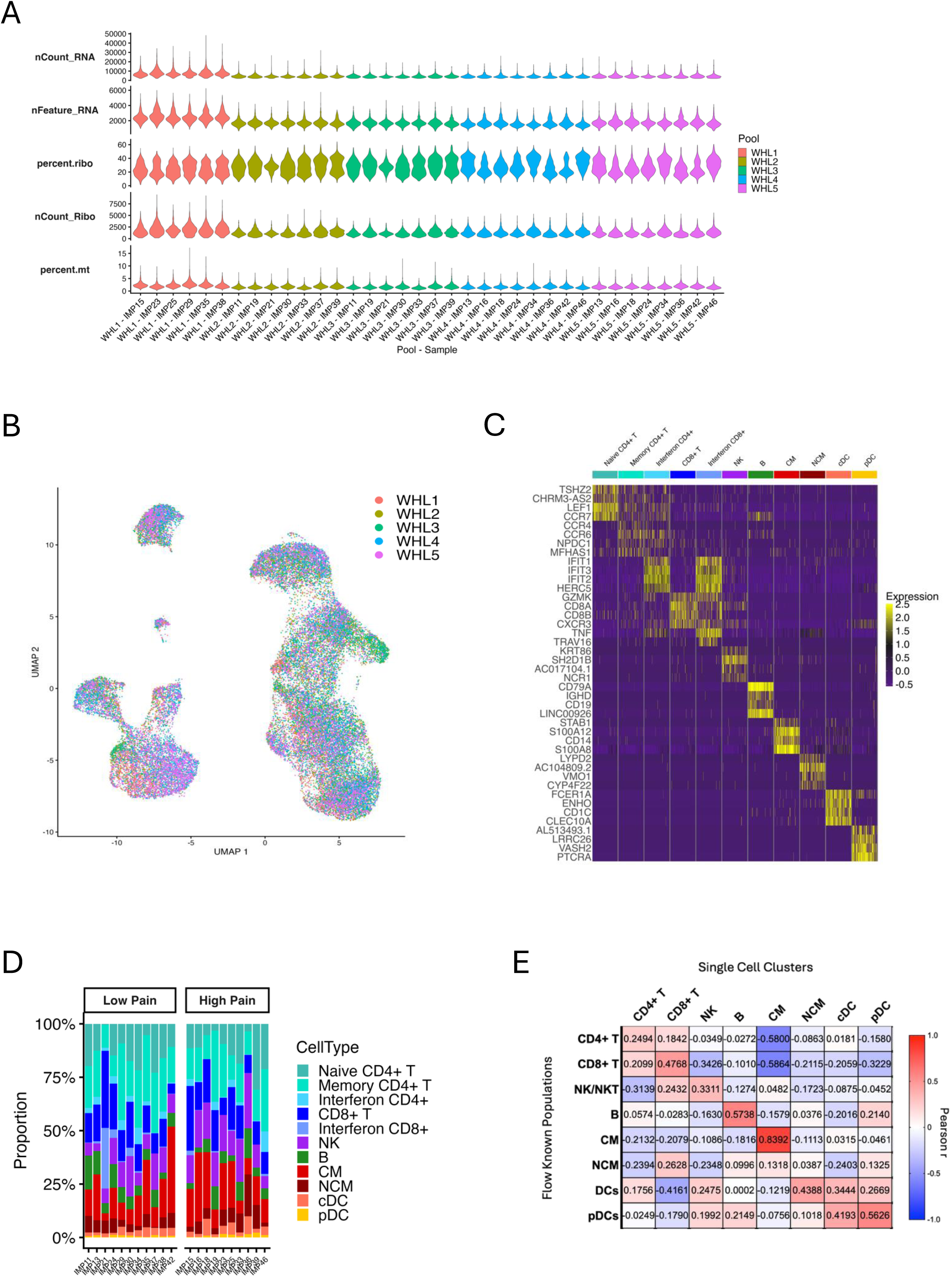
Additional single-cell RNA-seq analysis. (A) Violin plots of quality control metrics across samples with technical replicates grouped by sequencing pool. (B) UMAP annotated by sequencing pool. (C) Heatmap of top 4 *de novo* marker per cell population. (D) Stacked bar plot showing proportions of each cell population by sample grouped by pain. (E) Pearson correlation of cell populations proportions between flow and scRNA-seq analyses.

**Supp. Figure 4.**
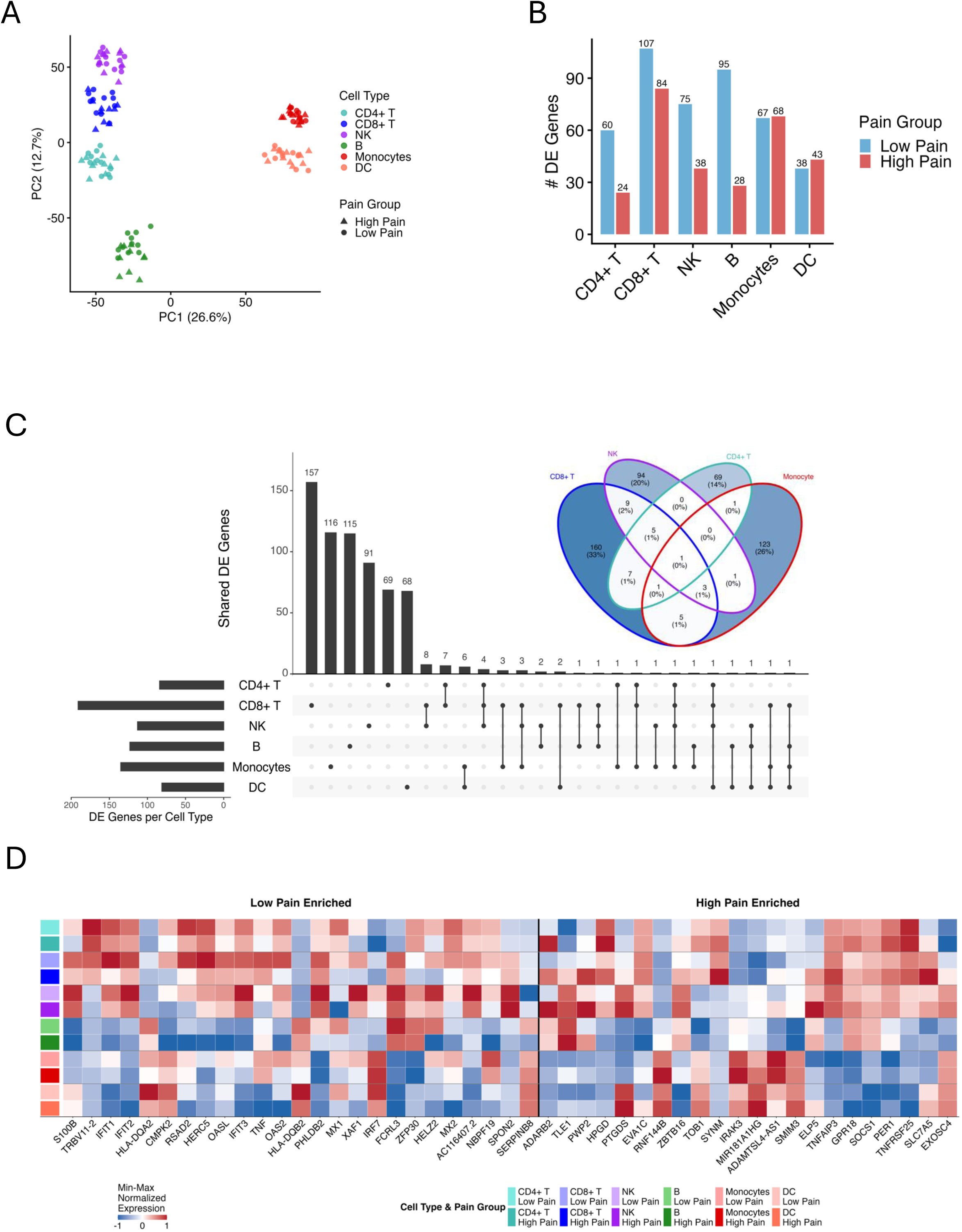
Additional pseudobulk differential expression figures. (A) PCA of global pseudobulk expression by cell types and sample from low (circle) and high (triangle) pain groups. (B) Bar plot showing the number of differentially expressed genes (DEGs, FC>|1.5|, p<0.05) by cell type associated with low (blue) or high (red) pain. (C) Upset plot and Venn Diagram quantifying DEG overlap between cell types. (D) Heatmap showing mean pseudobulk expression of DEGs that overlap between 1 or more cell types split by pain group.

